# A Simple Fluorescence Assay for Cystine Uptake via the xCT in Cells using Selenocystine and a Fluorescent Probe

**DOI:** 10.1101/2021.03.05.434035

**Authors:** Takashi Shimomura, Norio Hirakawa, Yuya Ohuchi, Munetaka Ishiyama, Masanobu Shiga, Yuichiro Ueno

**Author notes:** **Corresponding Author** Yuya Ohuchi – Dojindo Laboratories, Mashiki-machi, Kumamoto 861-2202, Japan.

## Abstract

The cystine/glutamate antiporter (xCT) is a crucial transporter that maintains cellular redox balance by regulating intracellular glutathione synthesis via cystine uptake. However, no robust and simple method to determine the cystine uptake activity of xCT is currently available. We have developed a method to measure the xCT activity via the reaction of selenocysteine and fluorescein *O,O′*-diacrylate (FOdA). Selenocystine, a cystine analog, is transported into cells through xCT on the cell membrane. The amount of the transported selenocystine was then determined by a reaction using tris(2-carboxyethyl)phosphine (TCEP) and FOdA in a weak acidic buffer at pH 6. Using this method, the cystine uptake activity of xCT in various cells, and the inhibitory efficiency of xCT inhibitors, were evaluated.

---

The cystine/glutamate antiporter (xCT) is an amino acid transporter that imports extracellular L-cystine with the exchange of intracellular L-glutamate across the cell membrane.^1^ The transported L-cystine is readily reduced to L-cysteine, and the L-cysteine is used as a substrate for the biosynthesis of glutathione (GSH), a major cellular antioxidant. Therefore, xCT is a critical transporter for the regulation of intracellular GSH levels to maintain cellular redox homeostasis.^2^ It has been reported that to escape from oxidative damage, several types of cancer cells overexpress xCTs, thus increasing intracellular GSH levels to scavenge reactive oxygen species (ROS), which is involved in resistance to cancer therapy.^3-5^ In glioma cells, excess levels of extracellular glutamate produced by highly expressed or up-regulated xCTs causes neurotoxicity implicated in neurodegenerative diseases.^6^ It has also been reported that cancer stem cells, which possess stem cell-like characteristics and contribute to tumor progression and metastasis, promote cystine uptake by stabilizing the xCT with a CD44 variant expressed on the cell surface.^7^ Thus, xCT plays an important role in maintaining tumor cell growth and survival, and is a potential target for cancer therapy. Recent studies have indicated that xCT inhibitors can exert an anti-tumor effect in cultured cancer cells and animal models, by disrupting cellular redox homeostasis and subsequent cell death.^8-12^ Ferroptosis, cell death mediated by the accumulation of iron-dependent lipid peroxides, can be induced by xCT inhibitors and has become of great interest in cancer therapy.^13-14^ Conventional cystine uptake assays require radioactive cystine analogs, such as ^14^C-cystine, ^3^H-cystine, or 35S-cystine,^15-18^ and special handling facilities. Thus, to avoid such restrictions and inconvenience, we have developed a novel and simple method to assess the cystine uptake activity of xCT in cells. This method uses selenocystine and fluorescein *O,O’*-diacrylate (FOdA). Selenocystine is a cystine analog that has selenium instead of sulfur. Makowske and Christensen have indicated that selenocystine can be taken up into cells through cystine transport-ers.19 Therefore, we used selenocystine to establish an xCT activi ty monitoring assay. To quantify the amount of selenocystine incorporated into the cells, we used FOdA. This fluorescent probe has been reported to react with cysteine and emit fluorescence^20^ and we predicted that selenocysteine would react with FOdA and emit fluorescence in the same manner.

The fluorescence intensities, and the spectra, of FOdA mixed with selenocystine and tris(2-carboxyethyl)phosphine (TCEP), and with only selenocystine or TCEP, are shown in Figures 1 and S1, respectively. As expected, strong fluorescence was observed in the solution after incubation of FOdA with both selenocystine and TCEP. In contrast, no fluorescence changes were observed in the other solutions. These results suggested that FOdA reacted with selenocysteine, a reduced form of selenocystine, and generated fluorescein as shown in the proposed mechanism in Figure 2. We also evaluated the reactivity of FOdA with selenocystine, cystine, and glutathione under various pH conditions in the presence of TCEP (Figure S2). All the solutions containing thiol compounds, and even the solution with no thiol compound (Blank), emitted fluorescence at pH 7. These results indicated that both the reaction with thiols and the hydrolysis of FOdA occurred at pH 7. However, it was found that a selective reaction of FOdA with selenocysteine proceeded at pH 6.5 or 6. Furthermore, cysteine and glutathione, the most abundant thiols in cells, did not interfere with the detection of selenocysteine (Figure S3). This specificity was caused by the difference in the p*K*a values of the thiols. The p*K*a values of selenocysteine, cysteine, and glutathione are 5.2, 8.3, and 8.7, respectively.^21,22^ Therefore, at pH values lower than 6.5, cysteine and glutathione cannot react with FOdA because of the protonation of the sulfur atom. Thus, we chose a weak acidic buffer at pH 6 for the selective detection of selenocystine using FOdA. The reaction between selenocystine and FOdA was effectively performed in MES buffer at pH 6 in a selenocystine-dependent manner (Figure S4).

**Figure 1.**
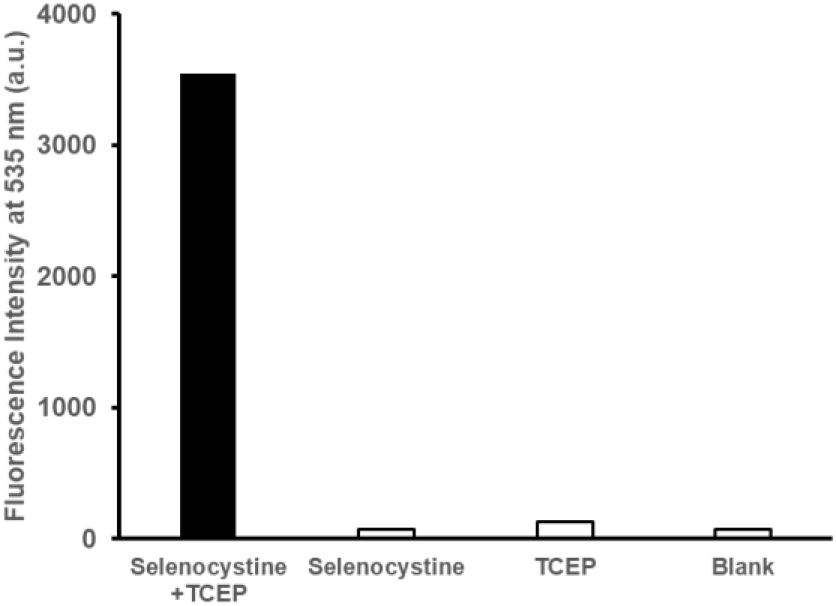
Fluorescence intensities at 535 nm of FOdA after reaction with selenocystine and TCEP. FOdA (10 µM) was incubated with or without selenocystine (10 µM) or TCEP (200 µM) at 37 °C for 30 min in 100 mM MES buffer (pH 6). The excitation wavelength was 470 nm.

**Figure 2.**
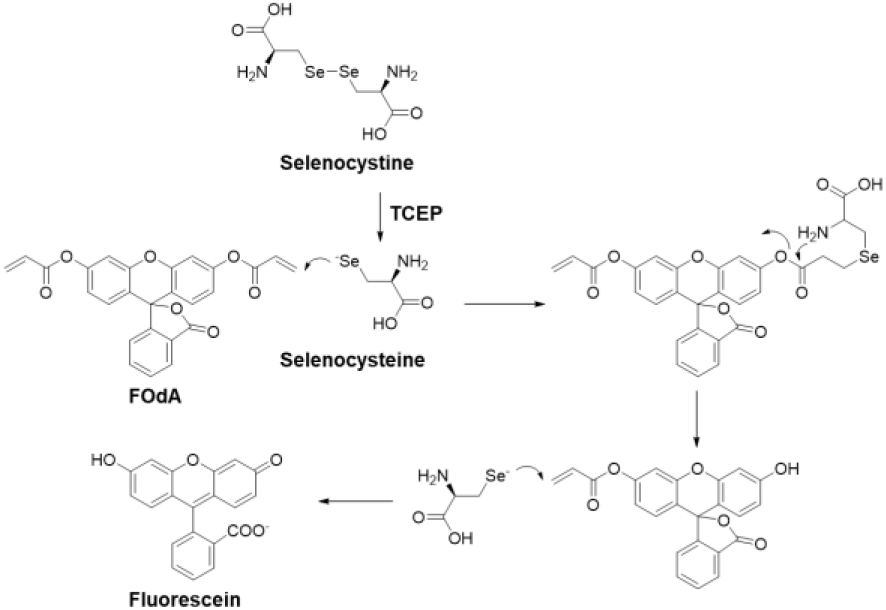
Possible reaction mechanism between FOdA and selenocystine in the presence of TCEP.

To evaluate whether selenocystine can be taken up into cells, HeLa cells were incubated with various concentrations of selenocystine, and the cell lysates were mixed with both FOdA and TCEP in 100 mM MES buffer at pH 6. As expected, the fluorescence increased depending on the concentration of selenocystine added to the cells (Figure S5). Moreover, this fluorescence response was effectively blocked by sulfasalazine and erastin, which are known xCT inhibitors (Figure 3). These results indicated that selenocystine was carried into the HeLa cells through the xCT on the cell membrane. The IC_50_ values of erastin and sul fasalazine toward xCT determined by this method were 0.054 and μM, respectively (Figure S6). Although these IC_50_ values cannot be directly compared with the values measured by the conventional method because both the cell types and experimental conditions are different, these data agree with a previous study in that the inhibition efficiency of erastin (IC_50_ = 1.4 μM) was much higher than that of sulfasalazine (IC_50_ = 26.1 μM).^23^ For further confirmation of the validity of this method, we used wild-type U251 glioblastoma cells that highly express xCT and xCT-knockout U251 cells.^23,24^ The wild-type U251 cells showed an intense fluorescence signal, but the xCT-knockout cells did not (Figure 6). These results strongly indicated that our method using selenocystine and FOdA worked well to determine cystine uptake activity, although the uptake rate of selenocystine might be different from that of cystine.

**Figure 3.**
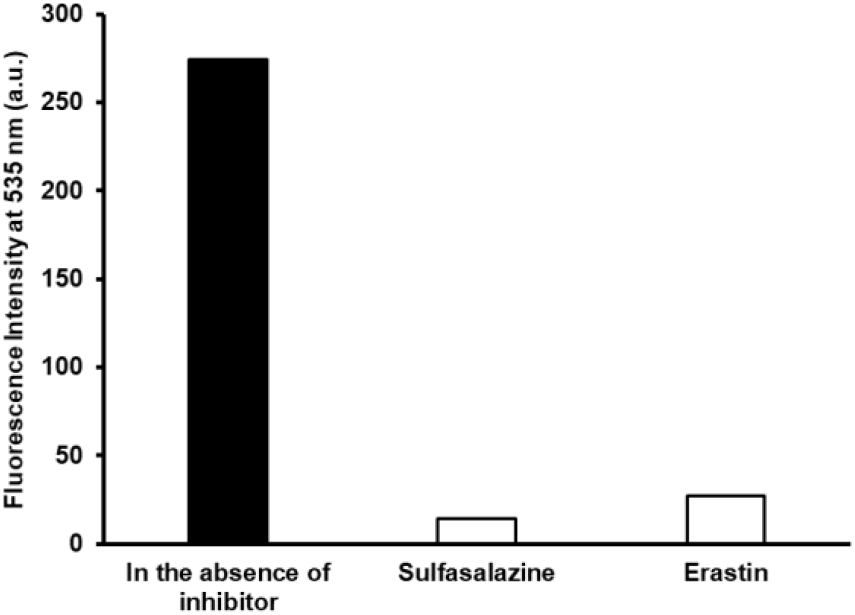
Fluorescence assay of cystine uptake activity in HeLa cells treated with xCT inhibitors (ex: 485 nm, em: 535 nm). The cells were incubated with selenocystine (200 µM) in the presence of sulfasalazine (0.5 mM) or erastin (2 µM), and then the cell lysates were reacted with FOdA (10 µM) in 100 mM MES (pH 6) containing TCEP (200 µM) at 37°C for 30 min.

**Figure 4.**
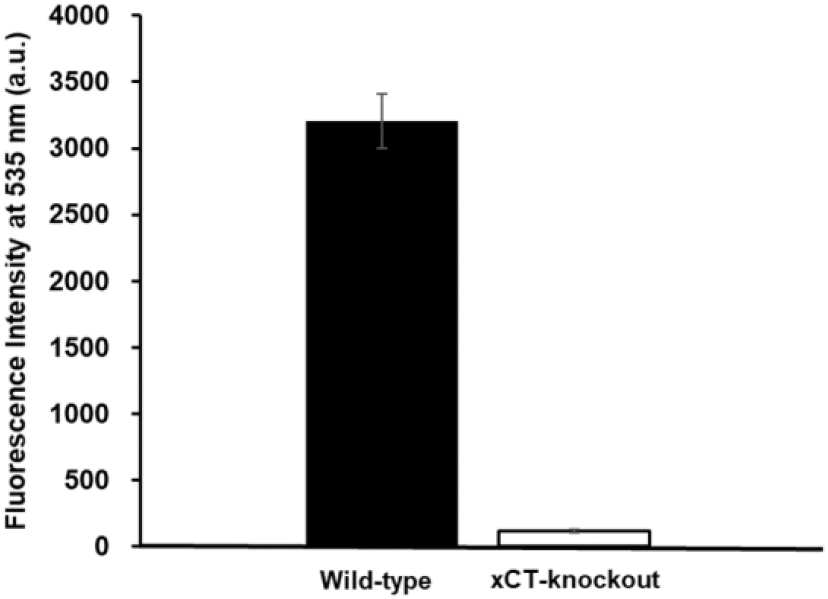
Fluorescence assay of cystine uptake activity in U251 wild-type or xCT-knockout cells (ex: 485 nm, em: 535 nm). The cells were incubated with selenocystine (200 µM), and then the cell lysates were reacted with FOdA (10 µM) in 100 mM MES (pH 6) containing TCEP (200 µM) at 37°C for 30 min.

Encouraged by these results, we performed a cystine uptake assay using HepG2 cells treated with diethyl maleate (DEM), which has been reported to induce xCT expression via the Keap1-Nrf2 system that acts as a defense against oxidative stress.^25^ After 24-h incubation of HepG2 cells in culture media containing various concentrations of DEM, the cystine uptake activity in the HepG2 cells was measured by the selenocystine and FOdA system. The results shown in Figure 5 indicated that the developed method can also be used to monitor changes in cystine uptake activity in response to oxidative stress levels.

**Figure 5.**
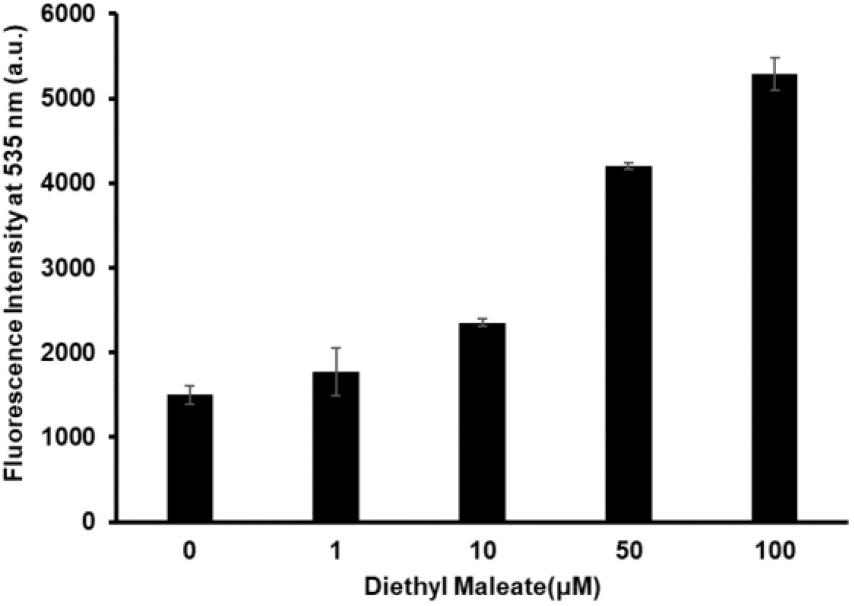
Fluorescence assay of cystine uptake activity in HepG2 cells treated with diethyl maleate (ex: 485 nm, em: 535 nm). The cells were treated with various concentrations of diethyl maleate for 24 h, followed by incubation with selenocystine (200 µM), and then the cell lysates were reacted with FOdA (10 µM) in 100 mM MES (pH 6) containing TCEP (200 µM) at 37°C for 30 min.

In conclusion, we have developed a safe, simple, and reliable method to measure cystine uptake activity using selenocystine as a cysteine analog, and FOdA as a fluorescent probe for selenocysteine, in combination with TCEP under weak acidic conditions. Because this method uses a simple process to measure the cysteine uptake activity, it will be possible to apply this method to microplate assays. Thus, this method will be useful to elucidate the regulation of cystine uptake associated with cellular metabolic alterations and oxidative stress in cancer cells and can be used in the discovery of potential anti-cancer drugs by high-throughput screening.

## ASSOCIATED CONTENT

### Supporting Information

The Supporting Information is available free of charge on the ACS Publications website at

Materials and methods (PDF)

## AUTHOR INFORMATION

### Authors

Takashi Shimomura − Dojindo Laboratories, Mashiki-machi, Kumamoto 861-2202, Japan; orcid.org/0000-0002-0215-2644

Norio Hirakawa − Dojindo Laboratories, Mashiki-machi, Kumamoto 861-2202, Japan

Munetaka Ishiyama − Dojindo Laboratories, Mashiki-machi, Kumamoto 861-2202, Japan

Masanobu Shiga − Dojindo Laboratories, Mashiki-machi, Kumamoto 861-2202, Japan

Yuichiro Ueno − Dojindo Laboratories, Mashiki-machi, Kumamoto 861-2202, Japan

### Notes

The authors declare no competing financial interests.

## ACKNOWLEDGMENT

We would like to thank Dr. Hironori Katoh (Kyoto University) for kindly providing both the U251 wild-type cells and xCT knockout cells.

## TOC graphic

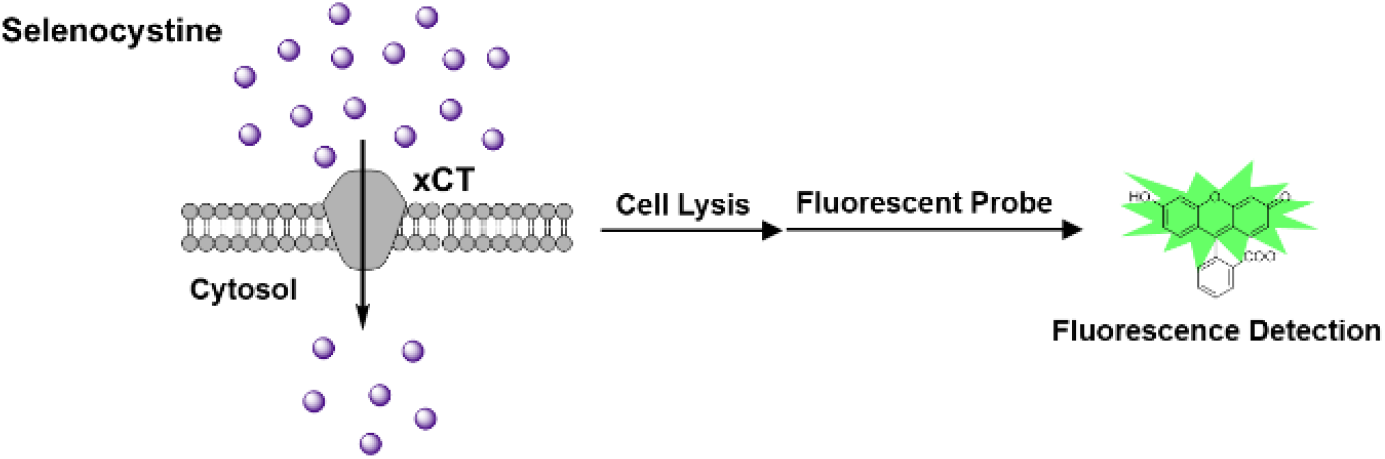

