## Supplementary material for "A Simple Fluorescence Assay for Cystine Uptake via the xCT in Cells using Selenocystine and a Fluorescent Probe"

**Table of contents**

**1. Materials and Methods**

**2. Supporting Figures**

### 1. Materials and Methods

#### General

Fluorescein *O,O'*-diacrylate (FOdA) was purchased from Sigma-Aldrich Chemical. Other reagents and solvents were purchased from FUJIFILM Wako Pure Chemical (Osaka, Japan) and Tokyo Kasei Industry (Tokyo, Japan). Minimum essential media (MEM), Dulbecco's Modified Eagle Medium (DMEM), Hank's balanced salt solution (HBSS), fetal bovine serum (FBS), non-essential amino acids, sodium pyruvate, L-glutamine, penicillin, and streptomycin were purchased from Thermo Fisher Scientific (Tokyo, Japan).

#### Fluorescence Measurements

Fluorescence spectra and intensities were measured by a microplate reader (Infinite M200PRO, TECAN). For fluorescence spectra measurement, the excitation wavelength was 470 nm. The fluorescence intensities at 535 nm were measured at an excitation wavelength of 485 nm. FOdA, which was dissolved in DMSO as a stock solution, was diluted with buffers to prepare 10 µM FOdA solutions. Measurements were performed before and after addition of thiol compounds and tris(2-carboxyethyl) phosphine hydrochloride (TCEP) to the FOdA solutions at final concentrations of 10 µM and 200 µM, respectively.

#### Cell Culture

HeLa cells were cultured in MEM with 10% FBS, 1% non-essential amino acids, 1 mM sodium pyruvate, 2 mM L-glutamine, 100 units/mL penicillin, and 100 µg/mL streptomycin.

U251 wild-type cells, U251 xCT-knockout cells and HepG2 cells were cultured in DMEM with 10% FBS, 4 mM L-glutamine, 100 units/mL penicillin, and 100 µg/mL streptomycin. All cells were incubated at 37 °C with 5% $CO_2$ in a humid atmosphere. To induce xCT expression, the HepG2 cells were cultured in the above medium containing various concentrations of diethyl maleate (DEM) under the same incubation conditions for 24 h.

#### Assay for Cystine Transporter Activity

Cell suspensions were added to a 96-well microplate at a total cell density of $1 \times 10^4$ cells/well and cultured at 37 °C with 5% $CO_2$ in a humid atmosphere overnight. After discarding the medium, the cells were incubated in a cystine- and serum-free medium at 37 °C with 5% $CO_2$ in a humid atmosphere for 5 min. The cells were washed with the cystine- and serum-free medium twice and incubated with a medium containing selenocystine under the same conditions for 30 min. After washing the cells with PBS three times, the cells were lysed with cold methanol and incubated with 100 mM MES buffer (pH 6.0) containing 10 µM FOdA and 200 µM TCEP at 37 °C for 30 min. Then, the fluorescence intensities were measured by a microplate reader (Infinite M200PRO, TECAN). The blank for the assay was measured by the same procedure but without using selenocystine.

### 2. Supporting Figures

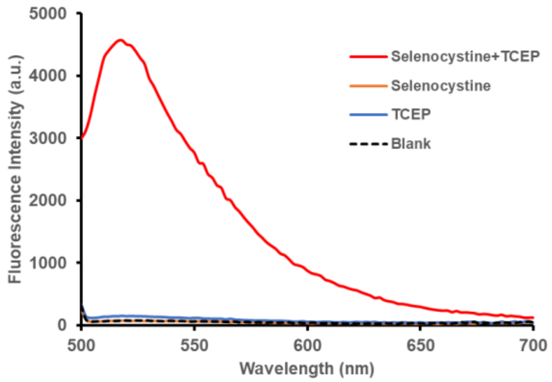


Figure S1. Fluorescence spectra of FOdA after reaction with selenocystine and TCEP. FOdA (10 µM) was incubated with selenocystine (10 µM) and TCEP (200 µM), and with only selenocystine (10 µM) or TCEP (200 µM) at 37 °C for 30 min in 100 mM MES buffer (pH 6). The excitation wavelength was 470 nm.

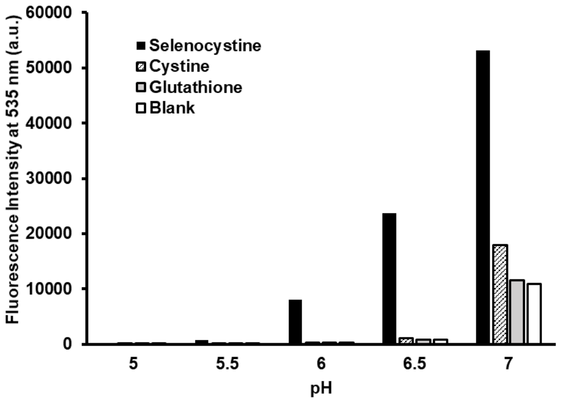


Figure S2. Fluorescence response of FOdA to selenocystine, cystine, and glutathione in the presence of TCEP in buffers of various pH values (ex: 485 nm, em: 535 nm). FOdA (10 µM) was incubated with selenocystine (10 µM), cystine (10 µM), or glutathione (20 µM) in the presence of TCEP (200 µM) at 37 °C for 30 min in 100 mM sodium acetate (pH 5 or 5.5), 100 mM MES (pH 6 or 6.5), or 100 mM sodium phosphate (pH7).

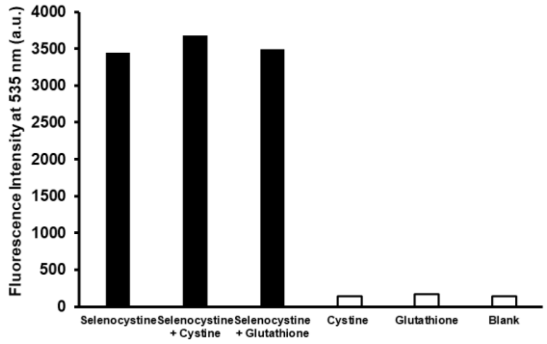


Figure S3. Fluorescence response of FOdA to selenocystine in the presence of cystine or glutathione (ex: 485 nm, em: 535 nm). FOdA (10 µM) was incubated with selenocystine (10 µM) in the presence or absence of cystine (10 µM) or glutathione (20 µM) in 100 mM MES (pH 6) containing TCEP (200 µM) at 37 °C for 30 min.

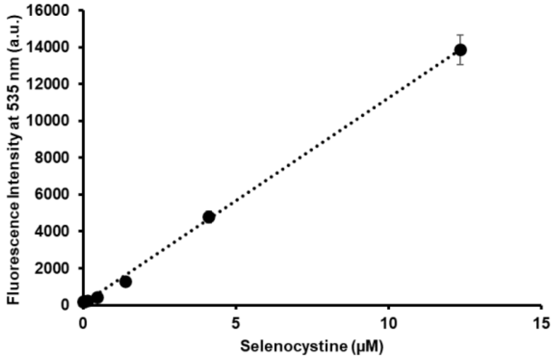


Figure S4. Fluorescence response of FOdA to various concentrations of selenocystine (ex: 485 nm, em: 535 nm). FOdA (10 µM) was incubated with different concentrations of selenocystine in 100 mM MES (pH 6) containing TCEP (200 µM) at 37 ℃ for 30 min.

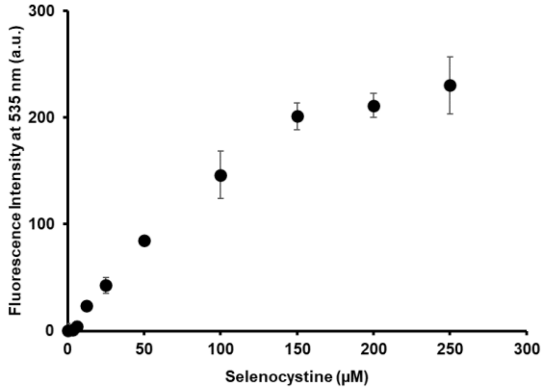


Figure S5. Fluorescence detection of selenocystine incorporated into HeLa cells (ex: 485 nm, em: 535 nm). The cells were treated with various concentrations of selenocystine, and then the cell lysates were reacted with FOdA (10 µM) in 100 mM MES (pH 6) containing TCEP (200 µM) at 37 ℃ for 30 min.

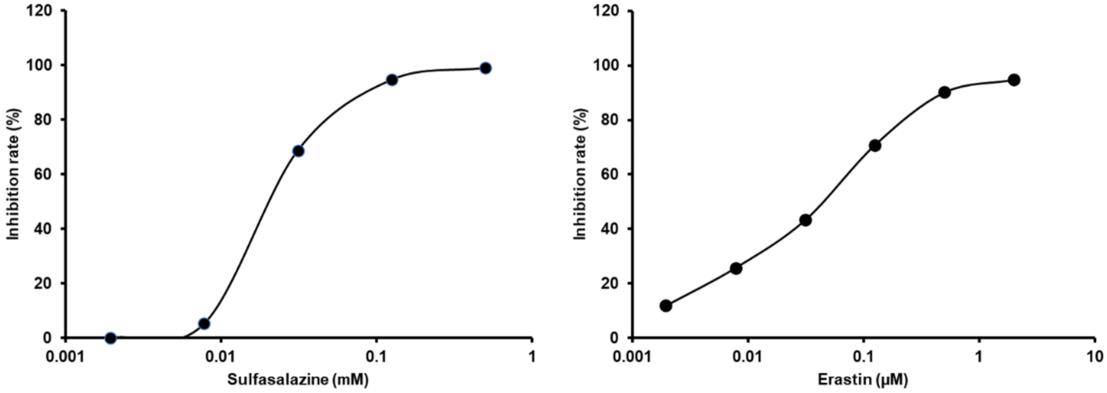


Figure S6. Inhibitory effect of xCT inhibitors on cystine uptake in HeLa cells. The cells were incubated with selenocystine (200 µM) in the presence of various concentrations of sulfasalazine or erastin, and then the cell lysates were reacted with FOdA (10 µM) in 100 mM MES (pH 6) containing TCEP (200 µM) at 37 ℃ for 30 min. The inhibition rates were calculated from the fluorescence intensities (ex: 485 nm, em: 535 nm) vs. the concentrations of the cystine transporter (xCT) inhibitors.
